# Differential expression of oncogenic lncRNAs NEAT1 and MALAT1 in 2D monolayer vs. 3D tumoroid culture and its implications in cancer progression

**DOI:** 10.1101/2024.10.22.619405

**Authors:** Arpita Ghosh, R. Soundharya, Mohit Jolly, Abhijit Majumder

## Abstract

Long non-coding RNAs (lncRNAs) have emerged as crucial regulators of cellular processes, overturning their previous classification as “junk” DNA. Their involvement in various cellular functions makes them potential therapeutic targets in a range of diseases, particularly in tumorigenesis and cancer. Among these lncRNAs, MALAT1 (Metastasis-Associated Lung Adenocarcinoma Transcript 1) and NEAT1 (Nuclear-Enriched Abundant Transcript 1) are prominent oncogenic lncRNAs with complex regulatory roles across multiple cancer types. Although the roles of these lncRNAs in cancer have been extensively studied, the majority of these investigations have been conducted in 2D culture systems. While 2D cell cultures are well-established and widely accepted models in cell biology, they lack the physiological complexity of 3D tumor architecture. In contrast, 3D cultures, where cells grow as three-dimensional clusters, better mimic *in vivo* conditions, making them essential for understanding tumor microenvironment (TME) dynamics. Despite the importance of 3D models, there is a lack of literature exploring lncRNA expression in 2D monolayers versus 3D cultures across different cancer types. This study examines the expression and function of the lncRNAs NEAT1 and MALAT1 in 3D tumoroids compared to 2D monolayer cultures, aiming to bridge the gap between *in vitro* models and the complex *in vivo* tumor microenvironment. We addressed this gap by quantifying the expression differences of NEAT1 and MALAT1 using qRT-PCR in breast cancer, liver cancer, cervical cancer, and glioblastoma (GBM) cells grown in both 2D and 3D cultures. Significant variations in NEAT1 and MALAT1 expression were observed between 2D and 3D cultures across these cancer types, signifying the need to study lncRNAs in 3D micro-environments. Furthermore, we established correlations between NEAT1 expression and cancer traits such as stemness, invasion, glucose transporter expression, and epithelial-mesenchymal transition (EMT) in GBM 3D tumoroids compared to 2D monolayers. Using siRNA to downregulate NEAT1 in GBM tumoroids, we demonstrated that reducing NEAT1 expression to the levels comparable to 2D cultures led to a decrease in the expression of mRNA markers associated with stemness, invasion, glucose transporters, and EMT. Additionally, siRNA-mediated downregulation of NEAT1 in 3D tumoroids directly impacted cancer properties, as validated by phenotypic assays, where reduced proliferation, migration, and invasion were observed in GBM (comparing 3D to 2D models). Therefore, our work provides new insights into the expression of two key oncogenic lncRNAs in 3D microenvironments of various cancers and lays the foundation for integrating lncRNAs as important molecular players within 3D culture systems, offering a better understanding of the *in vivo* complexity of the TME.

## Introduction

Cancer arises from mutations that disrupt the gene networks responsible for maintaining cellular balance. These alterations, involving both somatic and germline mutations, often occur in non-coding regions of the genome(1). As the name suggests, these regions instead of coding for proteins, are actively transcribed into non-coding RNAs (ncRNAs), with long non-coding RNAs (lncRNAs) being the most prominent (having length >200 base pairs)(2).

LncRNAs play diverse roles in cells, acting as scaffolds, guides, and decoys to regulate gene expression by forming complexes with DNA, RNA, or proteins(3, 4). They regulate key cellular processes: nuclear lncRNAs control chromatin and epigenetics, while cytoplasmic lncRNAs affect translation(3, 5, 6). They are vital for growth, development, and inflammation(7,8). In disease conditions such as cancer, lncRNAs exhibit dysregulated tissue-specific expression patterns and are involved in regulating cell cycle progression, survival, immune response, stemness maintenance, and pluripotency(3, 8–16). Recent studies have shown that lncRNAs may also engage in remodelling the tumor microenvironment and tumor metastasis(17, 18). Notable examples include NEAT1, which is a pro-oncogenic factor that plays a significant role in various solid tumors, such as liver, prostate, and gastric cancers, as well as renal cell carcinomas and glioma(19–26); MALAT1, which is widely associated with cancer and has been implicated in the regulation of alternative splicing and gene expression(27–33); HOTAIR, which promotes breast cancer metastasis by silencing key developmental genes(34); and ANRIL, associated with poor prognosis in prostate and gastric cancer by silencing the tumor suppressor locus(35, 36). These findings highlight the functional significance of lncRNAs in cancer and their potential as therapeutic targets. Despite the initial challenges in studying lncRNAs, advances in deep-sequencing technologies have significantly improved our understanding of their roles in cancer, revealing thousands of dysregulated lncRNAs across different cancer types.

Much of our understanding of lncRNAs comes from 2D monolayer cultures, but systematic studies in more physiologically relevant models are needed to clarify their roles in oncogenesis. Exploring lncRNA expression in the tumor microenvironment using 3D cell cultures offers advances over 2D systems(37-41). While 3D models better mimic *in vivo* conditions, providing insights into cell-cell interactions and tissue-like structures(42-44), the differences in lncRNA expression and function between 2D and 3D systems remain elusive. Despite significant research on gene expression in these models(45-50), studies on the non-coding RNA repertoire, particularly lncRNAs, are still unavailable. Exploring the changes in lncRNAs across 2D and 3D culture systems in turn will help identify which mRNAs or proteins may be affected by the lncRNAs, enhancing our understanding of disease progression at the molecular level and aiding in therapeutics or drug discovery. An existing study using a bioinformatics approach has indicated shifts in lncRNA profiles in breast cancer 3D vs. 2D cultures and identified dysregulated lncRNAs and their mRNA pairs(51). Yet, a comprehensive in-cellulo understanding of lncRNA dysregulation across diverse cancers in 3D versus 2D cultures or even the need to explore them is absent, potentially masking tissue-specific or universal lncRNA regulators impacting cancer pathways.

Our study aimed to bridge this gap by focusing on two well-studied oncogenic lncRNAs, MALAT1 and NEAT1, which are among the most upregulated lncRNAs pan-cancer(52–56). Utilizing TCGA (The Cancer Genome Atlas) correlation analysis, we established the clinical significance of MALAT1 and NEAT1 across Cervical, Liver, Breast, and GBM cancers. The qRT-PCR analysis in cell lines from these cancers cultured in 2D versus 3D tumoroids revealed varied upregulation of these lncRNAs in certain cancers in 3D models compared to 2D cultures, which instils the importance of understanding the tissue-specific expression patterns of lncRNAs in cancer. This study also shows that siRNA-mediated downregulation of NEAT1 in 3D GBM tumoroids led to significant reductions in proliferation, migration, and invasion. Notably, when NEAT1 levels in 3D tumoroids were downregulated to match those in 2D cultures, a marked decrease in cancer progression was observed, as supported by qRT-PCR analysis of invasion, stemness, and EMT markers. Functional assays further validated this reduction, demonstrating that the aggressive cancer properties seen in 3D tumoroids are closely linked to NEAT1 overexpression, a phenomenon not captured in 2D cultures for the same cancer cell line.

The study highlights the critical role of lncRNA expression variations in 2D and 3D cellular models, driven by distinct mechanical and micro-environmental factors. Understanding these tissue-specific expression patterns of lncRNAs is essential for developing targeted therapies, identifying unique drug targets, and enhancing diagnostic, prognostic, and therapeutic strategies for personalised and effective cancer treatments.

## Materials and method

### TCGA Correlation Analysis

TCGA data analysis was done using the “Correlation analysis” function in GEPIA 2(57). The correlation of the lncRNAs MALAT1 and NEAT1 with four pathways-Apoptosis, p53 and Epithelial Mesenchymal Transition and Proliferation, were estimated using Spearman correlation method. The signatures of the three pathways were obtained from the Hallmark gene set available on MSigDB. The colour coding was based on thresholding using the maximum and minimum values obtained for the Spearman correlation(58).

### 2D-Cell Culture

Cell lines such as HEK293 (embryonic kidney cells, obtained from Prof. V. Prasanna’s Lab, ACTREC, Navi Mumbai as a kind gift), HeLa (cervical cancer cells, obtained from Prof. Jyotsna Dhawan’s lab, CCMB Hyderabad as a kind gift), HepG2 (liver cancer cells, a kind gift from Prof. Jayesh Bellare’s Lab, Chemical Engineering Department, IIT Bombay), MCF7 (breast cancer cells, a kind gift from Prof. Dulal Panda, IIT Bombay), MDA-MB-231 (aggressive and invasive breast cancer cells, obtained from Prof. Sandip Kar’s Lab, Chemistry department, IIT Bombay as a kind gift), and U87-MG (aggressive glioblastoma cells, a kind gift from Prof. Shilpee Dutt’s Lab, ACTREC, Navi Mumbai) were utilized in this study. The cell lines underwent authentication, and their status was confirmed as mycoplasma-free. Maintenance involved culturing the cell lines in DMEM (Gibco) supplemented with 10% FBS (HiMedia), 10% antibiotic and anti-mycotic (Anti-Anti from Gibco), and 1% L-Glutamine (Gibco). Incubation of the cells were in a humidified incubator set at 37°C with 5% CO2. For 2D culturing, cells were seeded in 6-well plates at a density of 8×10^4^ cells per well and incubated for 48 hours or 72 hours according to the experiment performed. Subsequently, the cells were harvested for their respective experimental procedures.

### Generation of 3D-Tumoroids

For the generation of 3D tumoroids, a mimic of an ultra-low attachment plate was created by adding 50 µl of 2% agarose (prepared in serum-free DMEM media) to each well of a 96-well plate and allowing it to solidify. Then, 5000 cells from each cell line (as mentioned before) were added to each well. They were allowed to grow for 7 days in a humidified incubator at 37°C with 5% CO2. Media change was performed every alternate day or earlier if the media started to turn yellow. To achieve a satisfactory yield, 36 wells were seeded for tumoroids from each cell line for each biological replicate. For the NEAT1 knockdown tumoroids generated using the U87-MG GBM cell line, the cell suspension was first reverse transfected with siRNA to knockdown NEAT1 (methodology in the next section). Then, 15,000 cells were added to each well of the agarose-coated 96-well plate, where the cells were allowed to form tumoroids and grow for 72 hours. After this incubation period, the tumoroids were harvested for the respective experiments.

### Reverse transfection of siRNAs

Commercial ready-to-use siRNA targeting NEAT1 was purchased from Qiagen. The siRNA targeted the lncRNA between 0-1500 bp from the transcription start site. The transfection reagent Lipofectamine 3000 from ThermoFisher Scientific was used for the experiment. To generate NEAT1 downregulated 3D tumoroids with U87-MG cells, we used reverse transfection method to achieve maximum efficiency in siRNA-mediated knockdown. In reverse transfections, the transfection complexes are allowed to form within the wells before adding the cells. This method is quicker to perform than forward transfections and is preferred for high-throughput transfection applications. For transfection, an optimized concentration of 25 nM siRNA was used with 15,000 cells seeded in each well of a 96-well plate. The transfection was performed in complete media. The transfected cells were allowed to form tumoroids for 72 hours, after which they were harvested for respective experiments.

### RNA Extraction and cDNA Synthesis

RNA was isolated from the harvested 2D monolayer cells and 3D tumoroids using TRIZOL® reagent (Ambion®). The total RNA was isolated following the stepwise protocol as per the manufacturer’s instructions. cDNA was prepared from 1 µg RNA for each sample using a Biorad iScript™ cDNA synthesis kit according to the manufacturer’s instructions.

### qRT-PCR

Following cDNA preparation, real-time qPCR was conducted for all the samples with technical duplicates. The reaction volume for each was 15 µl, which included 1 µl of cDNA (after 1:2 dilution). The PCR conditions were as follows: Initial denaturation at 95°C for 3 minutes followed by 40 cycles of the following conditions:

95 °C for 10 s

60 °C for 30 s

72 °C for 30 s

qRT-PCR primers used in different experiments are listed in Supplementary Table 1. Transcripts were quantified using an iTaq™ Universal SYBR® Green Supermix (from Biorad) in the QuantStudio™ 5 Real-Time PCR instrument (Thermo). The Ct values obtained for the different transcripts were normalized to that of GAPDH. The fold change calculation for comparative analysis of the transcripts’ expression was done using the 2^-ΔΔCt^ method(59, 60). In brief, the fold change for each transcript was calculated using the following formula:

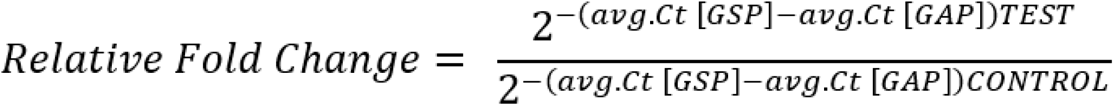

where avg. = average of; Ct = Cycle threshold; [GSP] = Gene Specific Primer; [GAP] = GAPDH Specific Primer; TEST and CONTROL are the respective experiment conditions.

### Ki67 Proliferation Assay

The U87-MG 2D monolayer cells and cells harvested from the 3D tumoroids were seeded at a density of 15,000 cells in a 12-well plate that already had 18 mm^2^ coverslips inserted in each well to which cells will be adhered. Post cell adherence to the coverslips, they were washed with 1x PBS. Cells on the coverslips from the wells were then fixed using a fixation buffer (3% paraformaldehyde, 5 mM EGTA [pH 8], 1 mM MgCl2) for 10 minutes. After fixation, cells were washed with washing buffer (30 mM glycine in PBS, 5 mM EGTA, and 10 mM MgCl2) twice, and then permeabilized using permeabilization buffer (0.2% Triton X-100 in PBS, 5 mM EGTA, and 10 mM MgCl2). After permeabilization, a similar washing step was followed. The wells containing the coverslips with adhered cells were next blocked using a blocking buffer (0.5% BSA in PBS, 5 mM EGTA, 10 mM MgCl2) for 30 minutes. After blocking was performed, the cells were incubated overnight with the primary antibody Ki67 (Thermo), which is a marker for proliferation, at a dilution of 1:300. Cells were washed thrice for 5 minutes each using blocking buffer post-incubation with the primary antibody. This was followed by incubation with the secondary antibody Alexa Fluor 488 (Invitrogen) at a 1:500 dilution for 2 hours. After incubation with the secondary antibody, cells were washed thrice for 5 minutes each using a blocking buffer. In each well, 600 mL of 1X PBS was added next, in which a Hoechst stain was added (1:3000 of 10mg/ml solution) and incubated for 10 minutes followed by a PBS wash. The coverslips were then viewed and analysed using an Evos FL Auto Microscope system under a 10X objective focus. Quantification for the images was done using ImageJ.

### 3D Migration and Invasion assay using Boyden’s chamber

Migration and invasion assays for the 2D monolayer cells and cells harvested from 3D tumoroids were conducted using Boyden’s chamber. In brief, 2D monolayer cells and cells from 3D tumoroids were counted and placed on the trans-well inserts at a density of 1.5×10^5^ cells for invasion and migration in FBS-free DMEM media and were allowed to migrate/invade for 24 hours at 37°C in a humidified incubator. Each well of the 24-well plate on which the trans-wells were placed to create the Boyden’s chamber already had complete media with FBS, acting as the chemo-attractant. The trans-wells for the 3D migration/invasion setup had a polycarbonate basement membrane that is permeable, and additionally, the trans-wells for invasion were coated with a layer of Matrigel (acting as an ECM) at a 1mg/ml concentration. Post 24-hour incubation, the inserts were taken out, non-migratory or non-invading cells from the top of the inserts were cleaned, and the cells that migrated/invaded through the polycarbonate basement membrane were fixed with methanol, stained using crystal violet dye, and images were taken using an EVOS FL microscope under the 10X objective. The dye from the stained cells on the basement membrane was extracted using methanol, and the optical density was recorded at 590 nm using a TECAN plate-reader and quantified.

### Statistics

Statistical analysis was conducted using GraphPad Prism 8.0 and Microsoft Excel to assess the significance across experimental replicates. Data are expressed as mean ± S.D. or ± S.E. (as specified in the figure legends) from three independent biological replicates, unless otherwise indicated. A two-tailed unpaired Student’s t-test was used for statistical evaluation, with a p-value < 0.05 considered statistically significant. Significance levels are indicated by one asterisk () for P < 0.05, two asterisks () for P < 0.01, three asterisks () for P < 0.001, and four asterisks (****) for P < 0.0001. The biological replicates used were consistent with standard practices in the field.

## Results

### Establishing MALAT1 and NEAT1 as clinically relevant oncogenic lncRNAs in various cancers using the TCGA database

The lncRNAs NEAT1 and MALAT1 are extensively studied due to their recognized oncogenic potential(28). NEAT1 has been implicated in various cancers, including breast, lung, colorectal, and glioblastoma, among others. Its dysregulation is associated with critical aspects of tumor behavior, such as growth, invasion, metastasis, and resistance to therapies, indicating its potential as both a biomarker and therapeutic target. NEAT1 primarily influences cancer progression by modulating gene expression and cellular pathways related to stemness, proliferation, apoptosis, epithelial-mesenchymal transition (EMT), and DNA damage response(61–66). Similarly, MALAT1 is linked to multiple cancers, exhibiting high expression levels that correlate with advanced tumor stages and poorer patient outcomes. It regulates genes associated with cell cycle regulation, DNA damage response, and EMT, contributing significantly to cancer progression and metastasis (67-71). Additionally, MALAT1 is associated with cancer stem cell-like properties, affecting tumor initiation, self-renewal, and resistance to therapy(67–71).

In our study, we evaluated the expression levels of NEAT1 and MALAT1 in 2D and 3D culture systems. To establish the clinical relevance of these two lncRNAs we assessed their correlation with hallmark cancer pathways (e.g., p53, apoptosis, EMT, and proliferation) (72, 73) using patient data from The Cancer Genome Atlas (TCGA) in four cancers: GBM, liver cancer, breast cancer, and cervical cancer. These four cancers were chosen based on literature reports indicating high levels of oncogenic MALAT1 and NEAT1 in these cancer types(52–56). Our analysis from TCGA revealed the maximum significant positive correlation of MALAT1 with P53, apoptosis, EMT, and proliferation pathways in liver cancer, while for NEAT1, the maximum significant positive correlation was observed with these pathways in GBM (Figure 1A (ii) and B (i), respectively). NEAT1 also showed a moderately positive correlation with these pathways in liver cancer (Figure 1B (ii)). In GBM, MALAT1 had a moderately strong positive correlation with the p53 and proliferation pathways, and a low positive correlation with the apoptosis and EMT pathways (Figure 1A (i)). There was a low positive correlation of MALAT1 with three out of the four hallmark cancer pathways in breast and cervical cancer (Figure 1A (iii, iv)). NEAT1 had a moderately positive correlation with two out of the four pathways in breast cancer (Figure 1B (iii)), and low positive, as well as negative, correlations with pathways in cervical cancer (Figure 1B (iv)). Although the pathway correlations of MALAT1 and NEAT1 in cervical cancer were the lowest among the four cancer types, various studies indicate the overexpression of MALAT1 and NEAT1 in cervical cancer and their involvement in related cancer pathways in disease progression(74–79). Thus, these findings, obtained from the TCGA data analysis and supported by existing literature, validate the clinical relevance for examining the expression and outcomes of the lncRNAs NEAT1 and MALAT1 in 3D tumoroids compared to their 2D cell monolayer culture.

**Figure 1.**
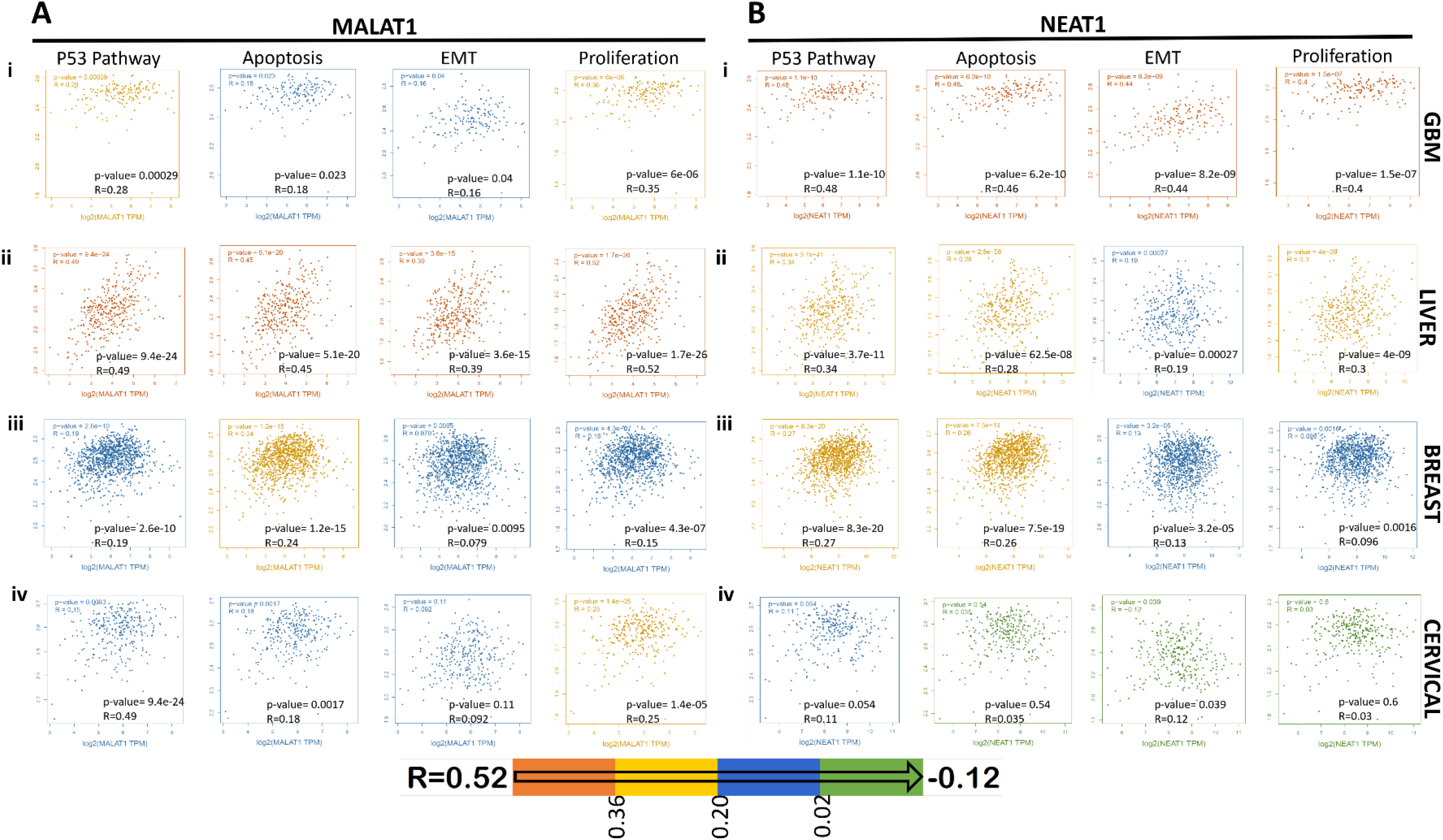
Establishing MALAT1 and NEAT1 as clinically relevant oncogenic lncRNAs in various cancers using the TCGA database. A(i-iv) Correlation of MALAT1 to cancer hallmark pathways in GBM, liver, breast and cervical tumors. B(i-iv) Correlation of NEAT1 to cancer hallmark pathways in GBM, liver, breast and cervical tumors. Colour coding below shows the trend from most to least positive correlation across the different cancer types.

### Differential expression of oncogenic lncRNAs MALAT1 and NEAT1 in 2D monolayer cells versus 3D tumoroids

After conducting TCGA correlation analyses for MALAT1 and NEAT1 across GBM, liver, breast and cervical cancer types, we aimed to assess the expression levels of these lncRNAs in cells cultured in 2D versus 3D tumoroids. The cell lines, HEK293 (normal kidney cells), HeLa (cervical cancer cells), HepG2 (liver cancer cells), MCF-7 (breast cancer cells), MDA-MB-231(aggressive breast cancer cels), and U87-MG (GBM cells) were cultured in 2D as outlined in the methods respectively. (Figure 2A(i) and B(i), Supplementary Figure S1A(i), B(i), C(i) and D(i)), and RNA was isolated after 48 hours. HEK293 human embryonic kidney cells served as the non-cancerous control to examine the lncRNA outcomes. Additionally, both MCF7 and MDA-MB-231 breast cancer cell lines were used to discern MALAT1 and NEAT1 expression differences between less and highly aggressive breast cancer forms. To generate 3D tumoroids or spheroids (for non-cancerous cells), an agarose-coated 96-well plate was employed (details in methods). Cells were cultured in these plates for 7 days to form tumoroids or spheroids (Figure 2A(ii) and B(ii) and Supplementary Figure S1A(ii), B(ii), C(ii) and D(ii)), which were then harvested for RNA isolation.

**Figure 2.**
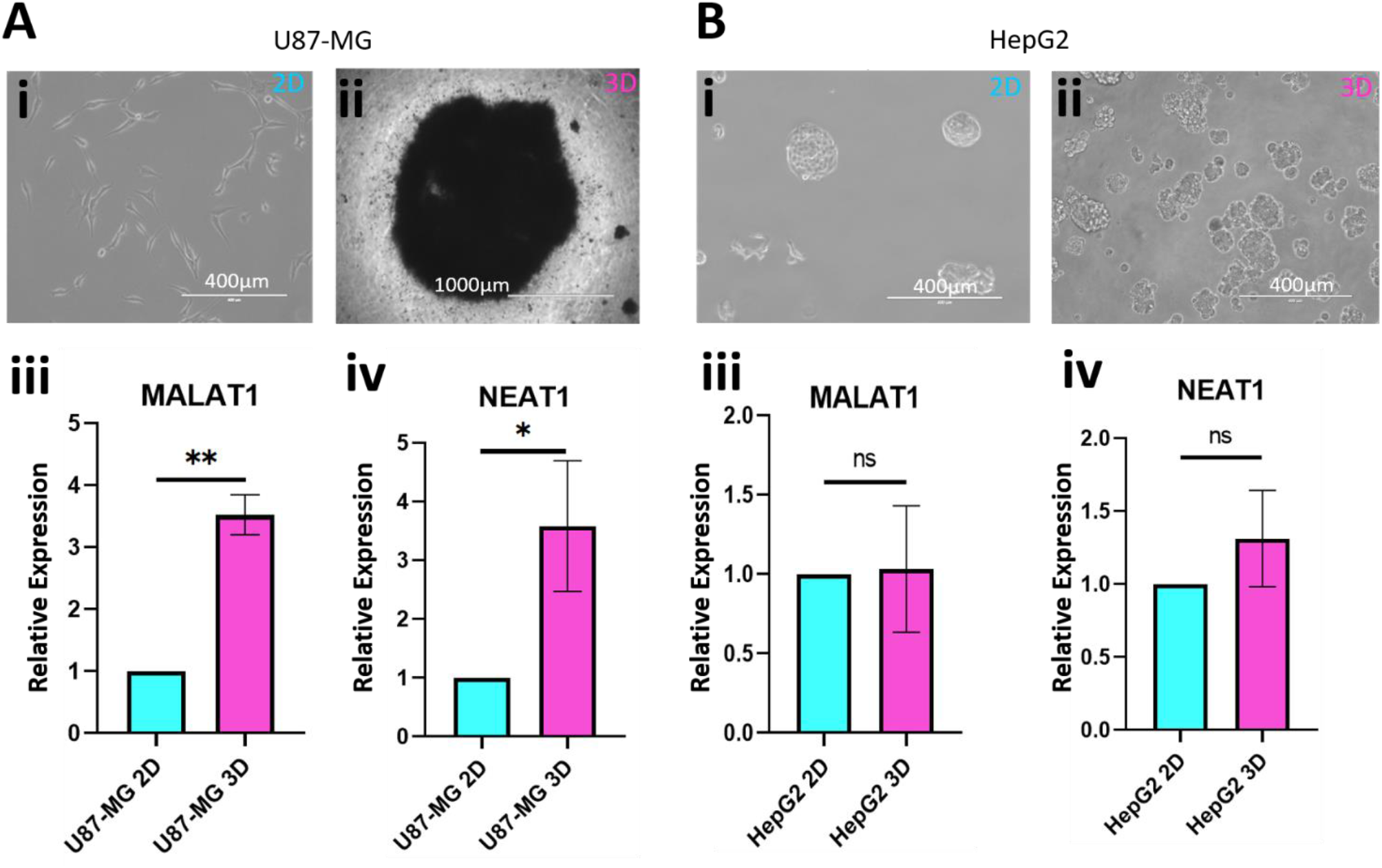
Differential expression of oncogenic lncRNAs MALAT1 and NEAT1 in 2D monolayer cells versus 3D tumoroids. (A) (i) 2D culture of U87-MG GBM cancer cell line. Scale bar: 400µm. (ii) 3D tumoroid observed after 7 days in U87-MG GBM cell line. Scale bar: 1000µm. (iii) Relative expression levels of lncRNA MALAT1 compared between U87-MG cells in 2D culture and 3D tumoroids. (iv) Relative expression levels of lncRNA NEAT1 compared between U87-MG cells in 2D culture and 3D tumoroids. (B) (i) 2D culture of HepG2 liver cancer cell line. Scale bar: 400µm. (ii) 3D tumoroid observed after 7 days in HepG2 liver cancer cell line. Scale bar: 400µm. (iii) Relative expression levels of lncRNA MALAT1 compared between HepG2 cells in 2D culture and 3D tumoroids in HepG2 cells. (iv) Relative expression levels of lncRNA NEAT1 compared between HepG2 cells in 2D culture and 3D tumoroids. Error bars in (A and B) represent ±S.D. across three independent biological replicates. *P<0.05, **P<0.01, ***P<0.001, and ****P<0.0001.

From the TCGA analysis, GBM and liver cancer showed the maximum positive correlation of NEAT1 and MALAT1 to the cancer hallmark pathways. Therefore, we first assessed the levels of MALAT1 and NEAT1 in 2D monolayer cultures versus the 3D tumoroids in GBM and liver cancer. Both MALAT1 and NEAT1 showed more than 3-fold upregulation in the 3D tumoroids as compared to 2D monolayer culture of GBM (Figure 2A (iii, iv)). Interestingly, no significant differential expression of NEAT1 and MALAT1 was found in HepG2 liver cancer monolayer cells and 3D tumoroids. (Figure 2B (iii, iv)). We also observed no significant differential expression of NEAT1 and MALAT1 in the spheroid culture of non-cancerous HEK293 as compared to their 2D monolayer culture (Supplementary Figure S1A(iii), (iv)). We further analysed the levels of MALAT1 and NEAT1 in the 2D monolayer cell cultures versus the 3D tumoroids of the breast (MCF7 and MDA-MB-231) and cervical cancers (HeLa). Both MALAT1 and NEAT1 showed up to 2.5-fold upregulation in the 3D tumoroids of MCF7 breast cancer cell lines. In MDA-MB-231 cell line MALAT1 showed 2-fold and NEAT1 showed 2.8-fold upregulation in the 3D-tumoroids (Supplementary Figure S1C (iii, iv) S1D (iii, iv)). Notably, NEAT1 was 12-fold upregulated in cervical cancer tumoroids vis-à-vis 2D monolayer culture, while MALAT1 did not show any significant difference (Supplementary Figure S1B (iii, iv)).

Therefore, to conclude, we found that in various cancer types, such as the breast cancer cell lines and GBM, both MALAT1 and NEAT1 were notably upregulated in 3D tumoroids. Conversely, in cervical cancer, one lncRNA is upregulated in 3D, while the other shows no significant change. Interestingly, in liver cancer, neither lncRNA displayed differential expression between 2D and 3D models. Also, the expression of oncogenic lncRNAs remained unaltered between 2D and 3D culture in non-cancerous HEK293 cells. Our observations highlight that in the different cancer cells from different tissue types, the expression of these lncRNAs may vary across 2D and 3D cultures. Hence, understanding these variations in lncRNA expression and selecting the most appropriate model that closely mimics the physiological scenario is crucial, as these lncRNAs play crucial roles in modulating downstream cancer regulatory pathways.

### Increased NEAT1 expression in 3D tumoroids compared to 2D monolayer culture increases cancer-related mRNA expression

From TCGA data analysis, we found that NEAT1 has maximum positive correlation to different cancer pathways in GBM (Figure 1B (i)). Also, our qRT-PCR studies show that for U87-MG cells NEAT1 expression was 3 to 4-fold upregulated in 3D tumoroids as compared to the 2D monolayer culture (Figure 2A (iv)). Next, we aimed to determine if U87-MG 3D tumoroids were more cancerous in nature than 2D monolayer cells and if this was linked to higher NEAT1 lncRNA expression in the 3D tumoroids. Therefore, we designed subsequent qRT-PCR experiments to evaluate gene expression levels from various cancer pathways which are linked to NEAT1 lncRNA such as stemness (OCT4, SOX2, NANOG, CD133), invasion (MMP2, MMP9, Fibronectin), glucose transporters (GLUT1, GLUT3), and EMT (Snail, Slug, Vimentin, N-Cadherin, Beta-catenin) (Supplementary Table 1). First, the lncRNA NEAT1 expression levels in 2D monolayer culture, 3D tumoroids, and NEAT1 downregulated 3D tumoroids were estimated (Figure 3A). To note, NEAT1 in the 3D tumoroids were downregulated to the level comparable to that in the cells from the 2D culture using an optimized concentration of NEAT1 targeting siRNAs as mentioned in the methods earlier. NEAT1 expression in 3D tumoroids was found to be 2.8-fold higher than in 2D monolayer culture. Upon downregulation of NEAT1 in the 3D tumoroids, its levels were reduced to match those observed in the 2D cells in monolayer (Figure 3A). We also simultaneously assessed whether the altered levels of NEAT1 in the cells affected the formation of these 3D tumoroids. However, we did not observe any differences in shape, size, or morphology between the control tumoroids and those formed from NEAT1-downregulated cells. (Supplementary Figure S2).

**Figure 3.**
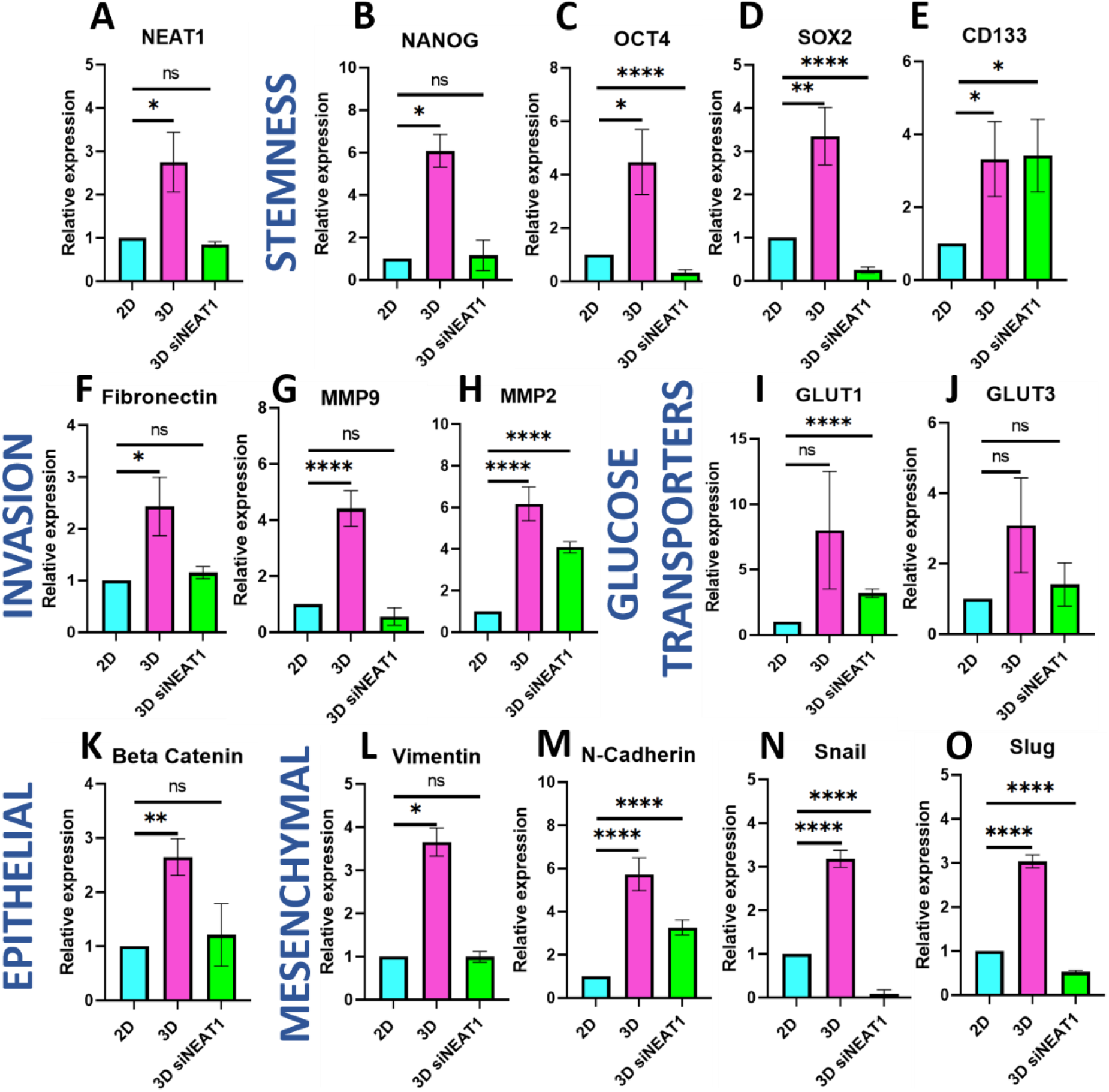
Increased NEAT1 expression in 3D tumoroids compared to 2D monolayer culture increases cancer-related mRNA expression. (A) Relative expression levels of NEAT1 in U87-MG cells grown in 2D culture, 3D tumoroids, and 3D tumoroids with siRNA mediated NEAT1 downregulation. (B-E) Relative expression of stemness markers (OCT4, SOX2, NANOG and CD133) in U87-MG cells grown in 2D culture, 3D tumoroids, and 3D tumoroids with siRNA mediated NEAT1 downregulation. (F-H) Relative expression of invasion markers (Fibronectin, MMP2 and MMP9) in U87-MG cells grown in 2D culture, 3D tumoroids, and 3D tumoroids with siRNA mediated NEAT1 downregulation. (I, J) Relative expression of glucose transporter (GLUT1 and GLUT3) in U87-MG cells grown in 2D culture, 3D tumoroids, and 3D tumoroids with siRNA mediated NEAT1 downregulation. (K) Relative expression of epithelial marker (Beta Catenin) in U87-MG cells grown in 2D culture, 3D tumoroids, and 3D tumoroids with siRNA mediated NEAT1 downregulation. (L-O) Relative expression of mesenchymal markers (Vimentin, N-Cadherin, Snail and Slug) in U87-MG cells grown in 2D culture, 3D tumoroids, and 3D tumoroids with siRNA mediated NEAT1 downregulation. Error bars in (A-O) represent ±S.D. across three independent biological replicates. *P<0.05, **P<0.01, ***P<0.001, and ****P<0.0001.

Next, the relative mRNA expression comparisons for the different cancer markers were made among the U87-MG cells from 2D culture, 3D tumoroids and NEAT1 downregulated 3D tumoroids. An elevated expression of stemness markers (OCT4, SOX2, NANOG, CD133) in 3D tumoroids versus 2D monolayer was observed, which upon downregulation of NEAT1 in the 3D tumoroids was mostly reversed (Figure 3B-E). NANOG, which exhibited a 6-fold upregulation in 3D tumoroids compared to 2D monolayer cells, showed expression levels similar to those in 2D monolayer cells in the NEAT1 downregulated 3D tumoroids (Figure 3B). On the other hand, CD133 stemness marker which had more than 3-fold upregulation in the 3D tumoroids compared to 2D monolayer culture, did not show any change in its expression upon NEAT1 downregulation in the 3D tumoroids (Figure 3E). The stemness markers OCT4 and SOX2 were upregulated by 4.5-fold and 3.5-fold, respectively, in 3D tumoroids compared to the 2D monolayer culture. However, upon NEAT1 downregulation in the 3D tumoroids, the expression of both markers significantly decreased, showing even more than 50% lower levels than those observed in the 2D monolayer culture. Likewise, invasion markers (MMP2, MMP9, Fibronectin), as well as glucose transporters (GLUT1, GLUT3) were prominently overexpressed in 3D tumoroids (Figure 3F-J). The invasion marker Fibronectin was upregulated approximately 2.5-fold in 3D tumoroids compared to 2D monolayer cultures. Upon downregulation of NEAT1 in 3D tumoroids, Fibronectin levels became comparable to those in 2D monolayer cells (Figure 3F). The invasion markers MMP9 and MMP2 also showed significantly higher expression in 3D tumoroids, with a 4-fold and 6-fold upregulation, respectively, compared to 2D cultures. However, in NEAT1 downregulated 3D tumoroids, MMP9 expression decreased to 50% lower than that in 2D cultures, while MMP2 expression reduced from 6-fold to 4-fold (Figure 3G, H). The expression of glucose transport markers GLUT1 and GLUT3 was upregulated in the 3D tumoroids, although it was not found to be statistically significant (Figure 3I, J). However, downregulation of NEAT1 in the 3D tumoroids caused a decrease in GLUT3 expression from 3 to 1.5-fold mRNA expression (Figure 3J) and the GLUT1 marker showed a stark downregulation, dropping from an 8-fold to a 3-fold mRNA expression (Figure 3I). Despite NEAT1 downregulation, both GLUT1 and GLUT3 expression levels in the 3D tumoroids remained higher than in the 2D monolayer cells. This observation was expected, as nutrient availability is limited in 3D tumoroids when compared to 2D monolayer cells, leading to an increased expression of these markers. Further, an epithelial marker (Beta-Catenin) and mesenchymal markers (Vimentin, N-Cadherin Snail and Slug) were upregulated in GBM 3D tumoroids compared to the 2D monolayer culture (Figure 3L-O). In NEAT1-downregulated 3D tumoroids, the expression of epithelial marker Beta-Catenin showed a significant decrease from 2.6-fold to 1.2-fold mRNA expression (Figure 3K). For mesenchymal markers, Vimentin expression in NEAT1-downregulated 3D tumoroids was comparable to that in 2D monolayer culture (Figure 3L). We found N-Cadherin mRNA levels dropping from 5.7-fold to 3-fold (Figure 3M). Interestingly, the mRNA levels of Snail and Slug were drastically reduced, by over 90% and 50%, respectively, compared to their expression in 2D monolayer cultures (Figure 3N, O).

Overall, these results demonstrate that NEAT1 significantly influences various aspects of GBM biology, including stemness, invasion, glucose metabolism, and EMT, with its effects more comprehensively reflected in 3D tumoroids compared to 2D cultures. This underscores the importance of studying lncRNAs like NEAT1 and its cancer-related pathways in 3D models in elucidating the complex interactions, offering valuable insights into tumor aggressiveness and potential therapeutic targets.

### Increased NEAT1 expression in GBM 3D tumoroids does not affect proliferation, but increases migration and invasion compared to 2D monolayer culture

As observed, the expression levels of NEAT1 lncRNA vary between GBM 3D tumoroids and 2D monolayer cultures, showing a direct correlation with key cancer-related properties such as stemness, invasion, and EMT. This correlation was confirmed through qRT-PCR analysis of specific markers. To determine whether these changes in mRNA expression also translate at the phenotypic level, we conducted functional assays to evaluate cell proliferation, invasion, and migration. These assays were performed on GBM cells grown in 2D culture, cells dissociated from 3D tumoroids, and cells from NEAT1 downregulated 3D tumoroids. A Ki67 assay was conducted to assess the proliferative potential of the cells under the different conditions(Figure 4A). U87-MG cells cultured in a 2D monolayer exhibited an average proliferation rate of 62% (Figure 4B). The cells dissociated from 3D tumoroids also had a comparable average proliferation rate of 58% (Figure 4B). However, the variation in proliferation rates observed for different cells was greater in the 3D tumoroids compared to those from the 2D monolayer culture. This increased variability may be due to the heterogeneity of cell populations within the 3D tumoroids with only its outer layer of cells being proliferative in nature. Upon downregulation of the NEAT1 lncRNA in the 3D tumoroids, the average percentage of proliferating cells dropped significantly to 28% with minimal variation.

**Figure 4.**
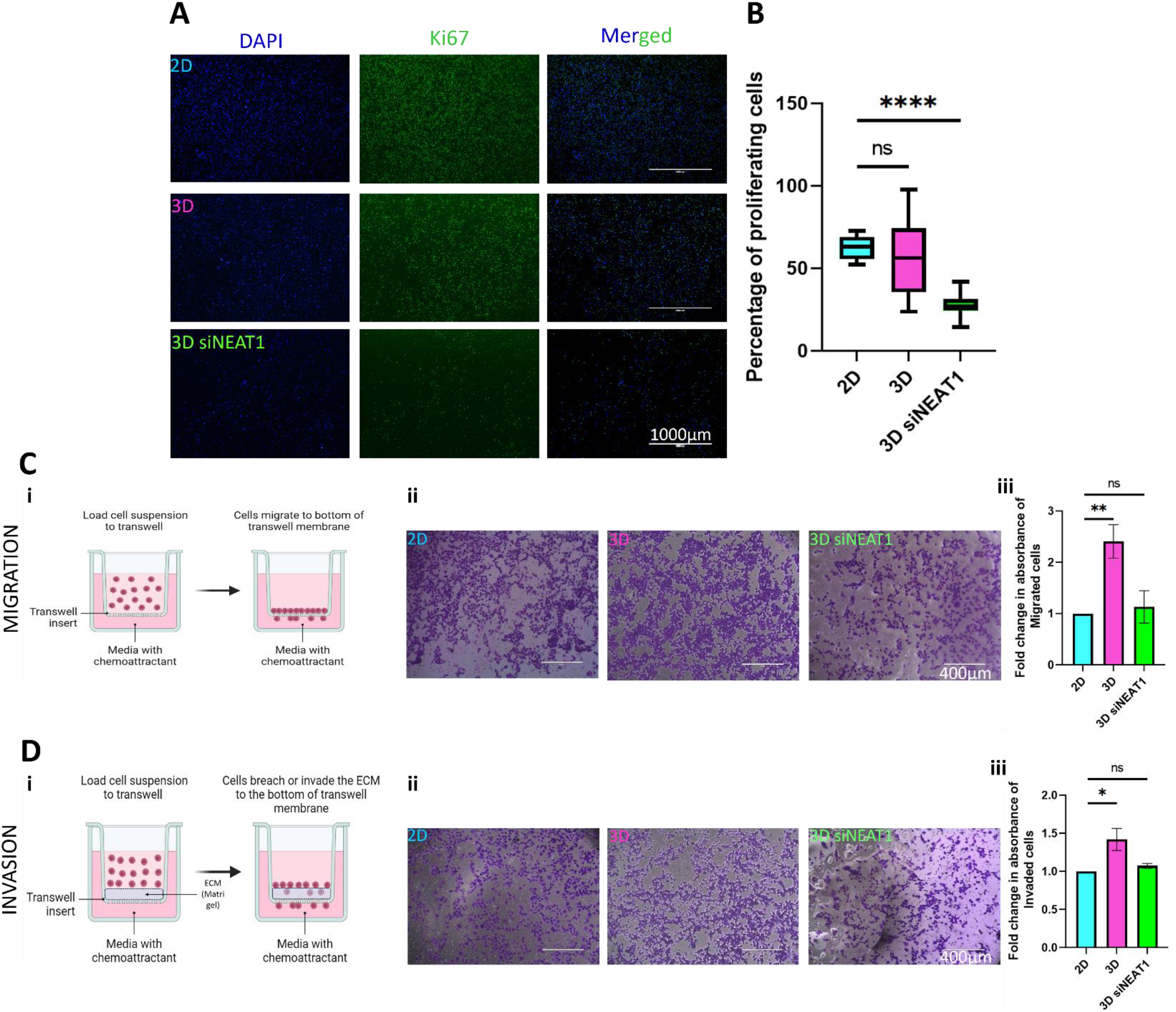
Increased NEAT1 expression in GBM 3D tumoroids does not affect proliferation, but increases migration and invasion compared to 2D monolayer culture. (A) Representation of Ki67 cell proliferation assay showing Ki67 positive cells as proof for the cells to be in the proliferative phase. The figure shows cells from 2D culture and dissociated cells from 3D tumoroids and NEAT1 downregulated 3D tumoroids. Scale bar corresponds to 400µm. (B) Quantification for the images from Ki67 proliferation assay is provided with error bars representing ±S.D.; **P<0.05, **P<0.01, ***P<0.001, and ****P<0.0001; Student’s t test. (C) Schematic representation to describe the process of (i) migration and (ii) invasion through the Boyden’s chamber. (D) Representation of cells from 2D culture and dissociated cells from 3D tumoroids and NEAT1 downregulated 3D tumoroids, which (i) migrated through the trans-well chambers (without Matrigel) and (ii) invaded through the Matrigel coated trans-well chambers. Scale bar corresponds to 400µm. (E) Fold change in the absorbance of the dye at 560 nm, extracted from the cells that (i) migrated or (i) invaded the trans-well chamber denoting the comparison in the motility and invasion properties of the cells from 2D culture and dissociated cells from 3D tumoroids and NEAT1 downregulated 3D tumoroids. Error bars represent ±S.E.; *P<0.05, **P<0.01, ***P<0.001, and ****P<0.0001.

To evaluate the migration and invasion potential of GBM cells grown under different conditions, we utilized the Boyden’s chamber, placing cells from both 2D monolayer and from 3D tumoroids on trans-well inserts, with invasion inserts coated in Matrigel. This approach was employed because invasion requires the cells to breach the extracellular matrix, in contrast to migration(Figure 4C). After 24 hours, migrated/invaded cells were stained with crystal violet, visualized, and quantified via optical density at 590 nm. Cells obtained from 3D tumoroids showed a 2.5-fold increase in migration potential compared to those from the 2D monolayer (Figure 4D(i) E(i)). However, migration potential of NEAT1 downregulated cells from 3D tumoroids was comparable to that of the 2D culture (Figure 4D(i) E(i)). For invasion, which requires cells to penetrate a Matrigel layer (i.e., an ECM layer) in the Boyden’s chamber (Figure 4C(ii)), we observed a significant but 0.5-fold increase in invading cells from 3D tumoroids compared to 2D monolayer cells (Figure 4D(ii) E(ii)). Downregulation of NEAT1 in 3D tumoroids resulted in invasion levels comparable to those of the 2D monolayer culture (Figure 4D(ii) E(ii)). These findings indicate that higher levels of NEAT1 in 3D tumoroids have a notable effect on the migration and invasion potential of GBM cells, with a more pronounced impact on migration.

Based on these observations, it can be concluded that elevated levels of NEAT1 in 3D tumoroids significantly affect the cancer properties of GBM cells compared to those in 2D monolayer culture. The increased variability in proliferation rates among cells from 3D tumoroids, coupled with the pronounced effect of NEAT1 downregulation on all proliferation, migration, and invasion processes underscores the crucial role of NEAT1 in modulating the aggressive behaviour of GBM cells in a 3D environment.

## Discussion

LncRNAs play crucial roles in cancer by regulating gene expression and influencing tumor progression, metastasis, and therapy resistance (4, 9, 18, 10–17). Despite their significance, the differential behavior of lncRNAs in 2D versus 3D cell cultures—more representative of *in vivo* conditions—remains underexplored. Understanding these differences is essential for developing accurate models for cancer research and therapy. Tissues in the human body exist in three-dimensional structures with unique cellular arrangements and stiffness. While 2D cell cultures are commonly used, they fail to replicate the complexity of physiological systems, contributing to numerous drug trial failures. This discrepancy is particularly critical in diseases like cancer, where solid tumors have distinct tumor microenvironments (TMEs) (37, 38). The formation of 3D tumoroids addresses aspects of the TME, preserving cellular arrangements and tumor stiffness that 2D systems cannot replicate (39, 40, 49, 50, 41–48). These tumoroids are widely used for drug screening due to their potential in identifying optimal drug candidates (80, 81). Research indicates substantial changes in the transcriptional and translational landscapes between 2D and 3D cell cultures (45, 46, 48–50). However, the shift in the profile of long non-coding RNAs (lncRNAs) between these environments remains relatively unexplored, with only one study addressing this. Nunez-Overa et al. (2022) investigated dysregulated lncRNAs in breast cancer by comparing 3D tumoroids and cells grown in 2D culture through microarray analysis (51), identifying 290 upregulated and 183 downregulated lncRNAs, and conducting co-expression analyses to explore oncogenic lncRNA-mRNA pairs. Their analysis revealed 102 lncRNA/mRNA pairs resembling those found in patient samples from TCGA datasets, but the findings were limited to bioinformatics and not validated *in vitro*.

In this study, we established the clinical relevance of two well-characterized oncogenic lncRNAs, NEAT1 and MALAT1, across multiple cancer types, including GBM, liver, breast, and cervical cancers (Figure 1), using TCGA correlation analysis. Our analyses confirmed positive correlations between NEAT1 and MALAT1 with pivotal cancer pathways such as p53, apoptosis, epithelial-mesenchymal transition (EMT), and proliferation across different cancers, with the strongest correlations observed in GBM and liver cancer. This observation laid the groundwork for studying the differential expression patterns of NEAT1 and MALAT1 in 2D versus 3D cancer tumoroids. We specifically focused on NEAT1 expression in GBM to demonstrate its impact on cancer progression in different cellular models.

We aimed to understand the differences in NEAT1 and MALAT1 expression in 2D monolayer culture versus 3D tumoroids for various cancer types by culturing GBM, breast, cervical, and liver cancer cells in both models (Figure 2, Supplementary Figure 1) and assessing lncRNA expression levels via qRT-PCR. This revealed intriguing variations: in breast cancer and GBM, both NEAT1 and MALAT1 showed elevated expression in 3D tumoroids, indicating their relevance to physiological contexts not captured in 2D cultures (Figure 2A and Supplementary Figure S1C, D). Conversely, in cervical cancer, NEAT1 was upregulated in 3D tumoroids, while MALAT1 levels remained unchanged (Supplementary Figure S1B). In liver cancer cells, no significant difference was observed in lncRNA expression between 2D and 3D culture models (Figure 2B), although NEAT1 and MALAT1 correlated strongly with hallmark cancer pathways in liver cancer according to TCGA datasets (Figure 1A(ii), B(ii)). This discrepancy may arise because HepG2 cells form 3D-like spherical colonies in 2D cultures, minimizing mechanical differences with their 3D tumoroids. Additionally, other liver-specific lncRNAs, apart from NEAT1 and MALAT1, might be dysregulated in sensing differences between these two culture systems, which requires further exploration. In the non-cancerous HEK293 cell model, lncRNA expression remained statistically insignificant between 2D and 3D cultures. Overall, we found that the distinct mechanical environments of cells in 2D cultures versus 3D tumoroids lead to varying expressions of these lncRNAs in the same cancer cells. The profiles of MALAT1 and NEAT1 highlight the tissue-specific nature of their differential expression, underscoring the need to explore more mechano-sensing lncRNAs or the cues controlling their expression for each cancer type.

The tissue-specific differences in lncRNA expression can be attributed to several factors. Micro-environmental differences play a significant role, as 2D cultures lack the complex interactions found in tissues. In contrast, 3D tumoroids better mimic the *in vivo* environment, affecting lncRNA expression. Mechanical cues also contribute, as 3D cultures replicate distinct tissue properties like stiffness and elasticity, leading to tissue-specific lncRNA expression patterns. The 3D environment can influence epigenetic modifications, such as DNA methylation and histone modifications, resulting in tissue-specific changes in lncRNA expression not observed in 2D cultures. Different tissues also have unique signaling pathways regulating lncRNA expression, with 3D cultures activating these pathways differently than 2D cultures. Lastly, tissue-specific transcription factors and co-factors may interact differently with lncRNAs in 3D versus 2D cultures, leading to variations in their expression (82, 83).

To explore the role of lncRNA expression in cancer progression between 2D and 3D culture models, we employed siRNA-mediated downregulation of NEAT1 in 3D GBM tumoroids, reducing its expression to levels comparable to 2D cultures. This approach allowed us to isolate the effects of NEAT1 overexpression due to the 3D cellular architecture. We found NEAT1 had the highest positive correlations with cancer progression pathways in clinical GBM samples, and expression levels differed between GBM 2D and 3D cultures. Thus, NEAT1 was chosen as a candidate lncRNA to study its effects on cancer progression in GBM models. Subsequent analysis of stemness, invasion, glucose transporters, and EMT markers in GBM cells grown in 2D, 3D, and NEAT1-downregulated 3D tumoroids, followed by functional assays, revealed the molecular dynamics driving increased cancer progression associated with altered lncRNA expression in 3D tumoroids.

When monitoring mRNA expressions of stemness markers in GBM cells grown in different conditions, we observed a significant increase in OCT4, SOX2, NANOG, and CD133 in the 3D tumoroids compared to 2D cultures (Figure 3B-E). However, NEAT1 downregulation in 3D tumoroids led to relatively unchanged CD133 levels, comparable NANOG levels to those in 2D cultures, and lower OCT4 and SOX2 expression than in 2D cultures. Earlier reports suggest that stemness in GBM relates to acquired drug resistance, leading to relapse and a more aggressive cancer (84, 85). Therefore, cancer stemness can be a critical parameter to study using an appropriate model system and this study suggests that three of the four tested stemness markers in 3D tumoroids are linked to NEAT1 levels, which vary greatly between 2D and 3D systems. With comparable NEAT1 levels in siRNA mediated NEAT1 downregulated 3D tumoroids and 2D monolayer cultures, we observed distinct differences in stemness properties, indicating that physiological scenario of GBM tumors could be better studied in 3D tumoroids.

Similarly, when studying mRNA expression of invasion markers, we observed that NEAT1 downregulation in 3D tumoroids affected Fibronectin, MMP2, and MMP9 (Figure 3F-H). All three markers were upregulated in 3D tumoroids compared to 2D cultures, but NEAT1 downregulation reduced MMP2 expression by ∼30%, although it remained higher than in 2D cultures. Fibronectin and MMP9 levels were much lower, comparable to or even lower than in 2D cultures. GLUT1 and GLUT3 glucose transporters were upregulated in 3D tumoroids compared to 2D cultures, although mRNA expression levels were not statistically significant (Figure 3I, J). NEAT1 downregulation in 3D tumoroids resulted in lower expression of these markers, although still higher than in 2D cultures, likely due to the unequal distribution of oxygen and nutrients in 3D tumoroids, which leads to higher receptor expression.

Furthermore, both epithelial and mesenchymal markers were highly expressed in 3D tumoroids, likely due to the heterogeneity of cellular populations within them. U87-MG GBM cells are E-Cadherin negative cell lines, and our conclusions stem from only one epithelial marker, Beta-Catenin, which was tested. Beta-Catenin was downregulated upon NEAT1 downregulation in 3D tumoroids, with expression levels comparable to 2D cultures (Figure 3K). Mesenchymal marker Vimentin followed a similar pattern to Beta-Catenin (Figure 3L), while N-Cadherin showed downregulation in mRNA expression in NEAT1 downregulated 3D tumoroids, yet the expression remained higher than in 2D cultures (Figure 3M). Interestingly, Snail and Slug were significantly downregulated with NEAT1 knockdown in 3D tumoroids compared to both 2D and 3D cultures.

Snail, a master regulator of EMT, plays a crucial role in cancer metastasis by repressing epithelial markers and promoting mesenchymal markers like N-Cadherin and Vimentin(86, 87). NEAT1, known to associate with Snail(88), differentially affects these EMT processes, particularly as seen here in the 3D tumoroids in GBM, highlighting the importance of critically analysing NEAT1’s impact on key cancer drivers.

The functional assays conducted in this study revealed that while the cellular proliferation rates in 2D and 3D cultures were comparable, there was a wide variation in the cells from 3D tumoroids (Figure 4A, B). NEAT1 downregulation in 3D tumoroids resulted in a stark decrease in proliferation compared to the cells both in 3D tumoroids and 2D cultures, with reduced variability (Figure 4B). It was intriguing to observe that the same levels of NEAT1 led to such significant differences in the proliferative rates of cells in 2D versus 3D cultures. Additionally, 3D tumoroids exhibited higher migration and invasion potential than 2D cultures, with a more pronounced effect of cell migration than invasion, but these levels were significantly reduced upon NEAT1 downregulation (Figure 4D, E), aligning with the observed mRNA marker expressions seen earlier. We hypothesize that the heterogeneity of cell populations in 3D tumoroids suggests that not all cells may be proliferating, with some acquiring EMT properties, as reflected in functional assays.

Overall, these findings underscore the differential roles of lncRNA NEAT1 in cancer progression across 2D and 3D culture systems, emphasizing the need for tailored research models to better understand lncRNA functions and their potential as therapeutic targets. Understanding lncRNA expression levels in 2D versus 3D systems is crucial for identifying unique drug targets and selecting appropriate platforms for drug screening, underscoring the importance of 3D models in cancer research. In conclusion, our study highlights the critical differences in lncRNA expression, specifically NEAT1 and MALAT1, between 2D monolayer cultures and 3D tumoroids across multiple cancer types, revealing their tissue-specific roles in cancer progression. These findings reinstate the importance of using 3D culture models to more accurately replicate the in vivo tumor microenvironment, thereby providing a more reliable platform for studying cancer biology and developing therapeutic interventions. The differential expression patterns observed suggest that lncRNAs like NEAT1 could serve as valuable biomarkers and potential therapeutic targets, offering new insights into cancer treatment strategies. Future research should focus on further elucidating the mechanisms behind lncRNA regulation in 3D models and exploring their implications in drug development.

## Acknowledgements

We deeply acknowledge the laboratories of Prof. Shilpee Dutt (ACTREC Navi Mumbai), Prof. V. Prasanna (ACTREC, Navi Mumbai), Prof. Jyotsna Dhawan (CCMB, Hyderabad), Prof. Dulal Panda (IIT Bombay), and, Prof. Jayesh Bellare (IIT Bombay) who kindly helped us by providing the desired cell lines for this study (as mentioned in the Methods section).

