## Supplementary Information for "Differential expression of oncogenic lncRNAs NEAT1 and MALAT1 in 2D monolayer vs. 3D tumoroid culture and its implications in cancer progression"

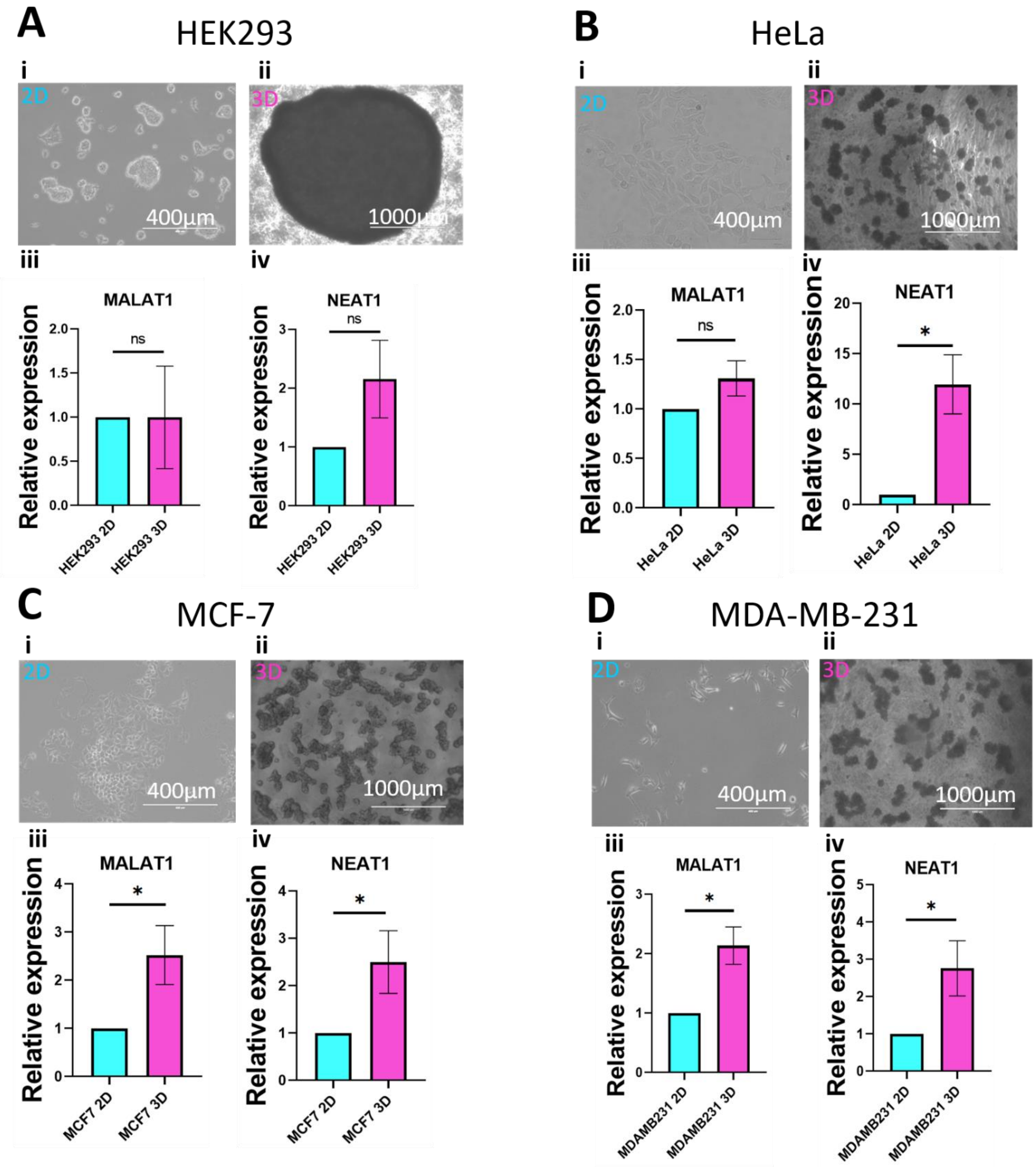


**Supplementary Figure S1. Differential expression of oncogenic lncRNAs MALAT1 and NEAT1 in 2D monolayer versus 3D tumoroids in different cancer cell types.** (A) (i) 2D culture of HEK293 non-cancer cell line. Scale bar: 400µm. (ii) 3D spheroid morphology observed after 7 days in HEK293 cell line. Scale bar: 1000µm. (iii) qRT-PCR to check relative expression levels of lncRNA MALAT1 compared between HEK293 cells in 2D culture and 3D spheroids. (iv) qRT-PCR to check relative expression levels of lncRNA NEAT1 compared between HEK293 cells in 2D monolayer cells and 3D spheroids. (B) (i) 2D culture of HeLa cervical cancer cell line. Scale bar: 400µm. (ii) 3D tumoroid observed after 7 days in HeLa cervical cancer cell line. Scale bar: 1000µm. (iii) qRT-PCR to check relative expression levels of lncRNA MALAT1 compared between HeLa cells in 2D culture and 3D tumoroids. (iv) qRT-PCR to check relative expression levels of lncRNA NEAT1 compared between HeLa cells in 2D culture and 3D tumoroids. (C) (i) 2D monolayer culture of MCF7 breast cancer cell line. Scale bar: 400µm. (ii) 3D tumoroid observed after 7 days in MCF7 breast cancer cell line. Scale bar: 1000µm. (iii) qRT-PCR to check relative expression levels of lncRNA MALAT1 compared between MCF7 cells in 2D culture and 3D tumoroids. (iv) qRT-PCR to check relative expression levels of lncRNA NEAT1 compared between MCF7 cells in 2D culture and 3D tumoroids. (D) (i) 2D monolayer culture of MDAMB-231 breast cancer cell line. Scale bar: 400µm. (ii) 3D tumoroid observed after 7 days in MDAMB-231 breast cancer cell line. Scale bar: 1000µm. (iii) qRT-PCR to check relative expression levels of lncRNA MALAT1 compared between MDAMB-231 cells in 2D culture and 3D tumoroids. (iv) qRT-PCR to check relative expression levels of lncRNA NEAT1 compared between MDAMB-231 cells in 2D culture and 3D tumoroids. Error bars in (A-D) represent ±S.D. across three independent biological replicates. *P<0.05, **P<0.01, ***P<0.001, and ****P<0.0001.

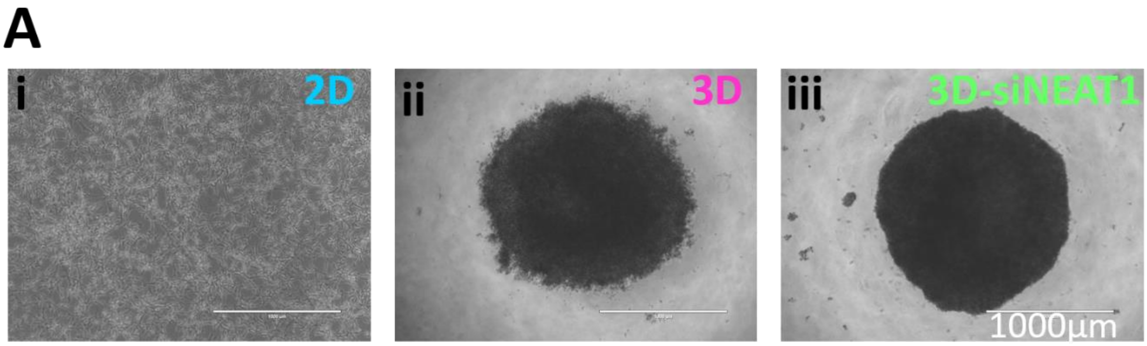


**Supplementary Figure S2. Microscopic image analysis to check the formation of 3D tumoroids post siRNA mediated NEAT1 downregulation.** (i) U87-MG cells in 2D monolayer culture grown for 72 hours and imaged before harvesting. Scale bar: 1000µm. (ii) 3D tumoroids formed with U87-MG cells for 72 hours as controls and imaged before harvesting. Scale bar: 1000µm. (iii) 3D tumoroids formed with NEAT1 siRNA reverse transfected cells for 72 hours and imaged before harvesting. Scale bar: 1000µm. Representative images are from three independent biological replicates (N=3).

**Supplementary Table 1. List of qPCR Primers**

| PRIMER NAME | SEQUENCE (5`-3`) |
|---|---|
| Vimentin FP | AGGCAAAGCAGGAGTCCACTGA |
| Vimentin RP | ATCTGGCGTTCCAGGGACTCAT |
| N-Cadherin FP | GCGTCTGTAGAGGCTTCTGG |
| N-Cadherin RP | GCCACTTGCCACTTTTCCTG |
| E-Cadherin FP | GGTTTTCTACAGCATCACCG |
| E-Cadherin RP | GCTTCCCCATTTGATGACAC |
| Beta Catenin FP | CACAAGCAGAGTGCTGAAGGTG |
| Beta Catenin RP | GATTCCTGAGAGTCCAAAGACAG |
| MALAT1 FP | GGGTGTTTACGTAGACCAGAACC |
| MALAT1 RP | CTTCCAAAAGCCTTCTGCCTTAG |
| NEAT1 FP | GCTGGACCTTTCATGTAACGGG |
| NEAT1 RP | TGAACTCTGCCGGTACAGGGAA |
| MMP2 FP | AGCGAGTGGATGCCGCCTTTAA |
| MMP2 RP | CATTCCAGGCATCTGCGATGAG |
| MMP9 FP | GCCACTACTGTGCCTTTGAGTC |
| MMP9 RP | CCCTCAGAGAATCGCCAGTACT |
| Fibronectin FP | ACAACACCGAGGTGACTGAGAC |

| | |
|---|---|
| Fibronectin RP | GGACACAACGATGCTTCCTGAG |
| CD133 FP | CACTACCAAGGACAAGGCGTTC |
| CD133 RP | CAACGCCTCTTTGGTCTCCTTG |
| Nanog FP | CTCCAACATCCTGAACCTCAGC |
| Nanog RP | CGTCACACCATTGCTATTCTTCG |
| Nestin FP | TCAAGATGTCCCTCAGCCTGGA |
| Nestin RP | AAGCTGAGGGAAGTCTTGGAGC |
| Glut1 FP | TTGCAGGCTTCTCCAACTGGAC |
| Glut1 RP | CAGAACCAGGAGCACAGTGAAG |
| Glut3 FP | CGTTGTTGGAATTCTGGTGGC |
| Glut3 RP | CTTAGCATTCTCCTCTTCTTTT |
| SOX2 FP | GCTACAGCATGATGCAGGACCA |
| SOX2 RP | TCTGCGAGCTGGTCATGGAGTT |
| OCT4 FP | CCTGAAGCAGAAGAGGATCACC |
| OCT4 RP | AAAGCGGCAGATGGTCGTTTGG |
| SLUG FP | ATCTGCGGCAAGGCGTTTTCCA |
| SLUG RP | GAGCCCTCAGATTTGACCTGTC |
| Snail FP | TGCCCTCAAGATGCACATCCGA |
| Snail RP | GGGACAGGAGAAGGGCTTCTC |
